# Atopic dermatitis complicated by recurrent eczema herpeticum is characterized by multiple, concurrent epidermal inflammatory endotypes

**DOI:** 10.1101/2023.02.27.530316

**Authors:** Nathan D. Jackson, Nathan Dyjack, Elena Goleva, Lianghua Bin, Michael T. Montgomery, Cydney Rios, Jamie L. Everman, Patricia Taylor, Caroline Bronchick, Brittany N. Richers, Donald Y. Leung, Max A. Seibold

## Abstract

**BACKGROUND:** A subgroup of atopic dermatitis (AD) patients suffer from recurrent, disseminated herpes simplex virus (HSV) skin infections, termed eczema herpeticum (EH), which can be life-threatening and contribute to AD morbidity. The pathobiology underlying ADEH is unknown.

**OBJECTIVE:** To determine transcriptional mechanisms of skin and immune system pathobiology that underlie ADEH disease.

**METHODS:** We performed whole transcriptome RNA-sequencing of non-lesional skin samples (epidermis, dermis) of AD patients with (ADEH^+^, n=15) and without (ADEH^-^, n=13) recurrent EH history, and healthy controls (HC, n=15). We also performed RNA-sequencing on plasmacytoid dendritic cells (pDCs) collected from these participants and infected *in vitro* with HSV-1. Differential expression, gene set enrichment, and endotyping analyses were performed.

**RESULTS:** ADEH^+^ disease was characterized by dysregulation in skin gene expression, which was limited in dermis (differentially expressed genes [DEGs]=14) and widespread in epidermis (DEGs=129). ADEH^+^-upregulated epidermal DEGs were enriched in type 2 cytokine (T2) (*IL4R, CCL22, CRLF2, IL7R*), interferon (*CXCL10, ICAM1, IFI44*, and *IRF7)*, and IL-36γ (*IL36G*) inflammatory pathway genes. At a person-level, all ADEH^+^ participants exhibited T2 and interferon endotypes and 87% were IL36G-high. In contrast, these endotypes were more variably expressed among ADEH^-^ participants. ADEH^+^ patient skin also exhibited dysregulation in epidermal differentiation complex (EDC) genes within the *LCE, S100*, and *SPRR* families, which are involved in skin barrier function, inflammation, and antimicrobial activities. pDC transcriptional responses to HSV-1 infection were not altered by ADEH status.

**CONCLUSIONS:** ADEH^+^ pathobiology is characterized by a unique, multi-faceted epidermal inflammation that accompanies dysregulation in the expression of EDC genes.

**Key Messages:**

1. AD patients with a history of recurrent EH exhibit molecular skin pathobiology that is similar in form, but more severe in degree, than in AD patients without this complication.
2. Non-lesional skin of ADEH^+^ patients concurrently exhibits excessive type 2 cytokine, interferon, and IL-36γ-driven epidermal inflammation.
3. Expression of these inflammatory skin endotypes among ADEH^+^ patients is associated with dysregulation in expression of epidermal differentiation complex genes involved in barrier function, inflammation, and antimicrobial activity.

**Capsule Summary:** AD patients with a history of recurrent disseminated HSV-1 skin infections form a unique molecular skin endotype group that concurrently exhibits type 2 cytokine, interferon, and IL-36γ-driven skin inflammation, accompanied by dysregulation in expression of epidermal differentiation complex genes involved in barrier function, inflammation, and antimicrobial activity.

## INTRODUCTION

Atopic dermatitis (AD) is a common chronic inflammatory disease of the skin, with a complex etiology driven by interacting genetic and environmental factors, which lead to both immune and skin barrier dysfunction. Despite these broad uniting factors, the clinical phenotype of AD is heterogeneous, with patients exhibiting different levels of severity, age of onset, symptoms, and duration of illness, as well as differing in their co-morbidities and disease complications.^1^

One important AD complication is increased susceptibility to bacterial and viral skin infections.^2, 3^ Aside from *Staphylococcus aureus* (*S. aureus*), the most common pathogen for AD skin is herpes simplex virus 1 (HSV-1),^2, 3^ which can lead to disseminated HSV-1 infection, termed eczema herpeticum (EH).^4^ EH is a devastating exacerbation of AD that requires immediate antiviral therapy. The etiology of EH is an epidemiological quandary: while 62% of adults are HSV-1 seropositive, only 3% of AD patients develop EH-like symptoms.^5, 6^ Moreover, there is a remarkable bimodality in the ‘recurrence’ of HSV infection; most individuals have only a single episode, but a small subgroup of AD patients has recurrent episodes (ADEH^+^ patients). The discrepancy between probable HSV-1 exposure and EH emergence suggests intrinsic host factors may determine the risk for EH development, and that those with ADEH^+^ disease may exhibit unique AD pathobiology.

Multiple studies support intrinsic differences in the pathobiology of ADEH^+^ patients compared to patients without a history of EH (ADEH^-^). For example, studies have found that ADEH^+^ patients have more severe disease, with early age disease onset, more frequent history of other atopic disorders, and high rate of skin *S. aureus* infections^4^. Additionally, ADEH^+^ patients have been shown to exhibit higher frequencies of both *FLG, TSLP and IL7R* genetic risk variants, compared to ADEH^-^ patients.^5, 7, 8^ A study of peripheral blood mononuclear cells (PBMCs) and CD8^+^ T-cells from ADEH^+^ individuals also found diminished capacity to produce IFN-γ in response to HSV-1 compared to ADEH^-^ individuals, suggesting systemic immune responses may differ in ADEH^+^ patients.^9^

Few studies have evaluated skin dysregulation in ADEH^+^ patients, but the occurrence of EH regardless of humoral immunity to HSV, suggests that skin innate immunity may play a critical role in ADEH^+^ susceptibility. Importantly, type 2 cytokine (T2) inflammation is a strong regulator of innate immunity, and biomarkers of T2 inflammation (blood eosinophils and serum IgE) have been found to be upregulated in ADEH^+^ patients^4^. Multiple skin transcriptomic studies of AD have documented different inflammatory AD endotypes, including those driven by type 2, type 1, type 17, and IL-36γ cytokines.^1, 10-14^

Given the unclear etiological relationship between AD and EH, we sought to characterize molecular differences between ADEH^+^ and ADEH^-^ skin based on transcriptomic profiling of non-involved (non-lesional) skin biopsies from these patient groups matched by disease severity. In doing so, we assayed both epidermal tissue, the primary origin of barrier function in the skin, and dermal tissue, wherein many tissue-resident immune cells reside. We also examined HSV-1-stimulated blood-circulating plasmacytoid dendritic cells (pDCs) from these individuals, which are involved in antiviral host responses in the skin, and may underlie differences in susceptibility to infection between ADEH^+^ and ADEH^-^ individuals. Based on these data, we define the distinct inflammatory pathways that underlie ADEH^+^ disease.

## METHODS

### Human subject recruitment

Skin samples were obtained from 15 adult AD patients with history of EH (ADEH^+^), 13 adult AD patients with no history of EH (ADEH^-^), and 13 nonatopic healthy control (HC) subjects. Eczema Area and Severity Index (EASI), Scoring Atopic Dermatitis (SCORAD), and Rajka-Langeland total scores were recorded, and serum IgE levels were measured (Table 1). All ADEH^+^ patients had prior history of ≥3 episodes of EH.

**Table 1.** Clinical characteristics of the study subjects.

| **Characteristic** | **ADEH<sup>+</sup>**<br>**n=15** | **ADEH<sup>-</sup>**<br>**n=13** | **HC**<br>**n=13** | **p.**<br>**overall** |
| --- | --- | --- | --- | --- |
| **Age (years)** |  |  |  |  |
| Mean (SD) | 33.1 (10.5) | 37.7 (11.4) | 41.0 (8.4) | 0.134 |
| **Gender, n (%)** |  |  |  |  |
| Female | 9 (60.0) | 9 (69.2) | 10 (76.9) | NA |
| Male | 6 (40.0) | 4 (30.8) | 3 (23.1) |  |
| **Ethnicity, n (%)** |  |  |  |  |
| Hispanic or Latino | 1 (6.7) | 1 (7.7) | 1 (7.7) | NA |
| Not Hispanic or Latino | 14 (93.3) | 12 (92.3) | 12 (92.3) |  |
| **Race, n (%)** |  |  |  |  |
| Caucasian | 13 (86.7) | 11 (84.6) | 10 (76.9) | NA |
| Black or African American | 2 (13.3) | 2 (15.4) | 3 (23.1) |  |
| **EASI Score** |  |  |  |  |
| Mean (SD) | 17.5 (11.0) | 15.9 (14.3) | NA | 0.749 |
| **SCORAD** |  |  |  |  |
| Mean (SD) | 50.8 (13.8) | 43.1 (18.6) | NA | 0.224 |
| **Rajka-Langeland Total**<br>**Score** |  |  |  |  |
| Mean (SD) | 7.3 (1.2) | 6.9 (1.8) | NA | 0.459 |
| **Serum IgE**<br>Median [1<sup>st</sup>; 3<sup>rd</sup> quartile] | 1100 [66;<br>2692]* | 300 [91.7;<br>841]** | 29 [7.5;<br>44.5] | 0.0002 |
| **Skin *Staphylococcus aureus* culture positive, n (%)** | 11 (73.3) | 8 (61.5) | 0 (0) | 0.0002 |

NA – not applicable.
\*p<0.001 as compared to NA, \*\*p<0.05 as compared to NA

### Skin biopsy collection and RNA extraction

Two-mm skin punch biopsies were performed in non-lesional skin of the upper extremities. Biopsy specimens were dissected into the epidermis and dermis, lysed, and RNA was isolated using RNeasy Micro Kits (Qiagen).

### Peripheral blood pDCs isolation and HSV stimulation

Heparinized venous peripheral blood was collected from each study participant. PBMCs were isolated and pDCs were purified using EasySep™ Human Plasmacytoid DC Isolation kits (STEMCELL™ Technologies). pDCs were mock- or HSV-1-treated for 20 hours, followed by RNA extraction.

### Keratinocyte culture and stimulation

Normal human embryonic keratinocytes (NHEK) were differentiated in culture for 3 days, then treated with IL-36γ (R&D systems) for 2 days, after which cells were harvested for RNA extraction. RNA-seq analysis and qRT-PCR were performed.

### RNA transcriptome gene expression and quality control

RNA-seq libraries were constructed and barcoded using the Ion AmpliSeq™ Transcriptome Human Gene Expression Kit. Barcoded RNA-seq libraries were pooled and sequenced on the Ion Torrent Proton sequencer.

### Analysis of bulk RNA-seq data

Unpaired single-gene differential analysis was carried out using the *DESeq2* R package^15^. For differential expression analysis between paired mock- and HSV-1-stimulated pDC samples, we used the *lmerSeq* R package based on variance stabilized transformation (VST)-normalization counts. We adjusted p-values to control for false discovery rate (FDR) using the Benjamini-Hochberg method. Functional enrichment analyses were performed using the EnrichR API^16^, weighted gene co-expression network analysis (WGCNA)^17^ was carried out using the *WGCNA* R package^18^, and gene set enrichment analyses (GSEA), were performed using GSEA v4.1.0^19^.

For additional details concerning study subjects, experimental procedures, transcriptomics data mapping, quantification, quality control, and data analysis, see Supplementary Methods.

## RESULTS

### ADEH^+^ patients exhibit more severe dysregulation of common AD skin disease pathobiology

To identify ADEH disease mechanisms, we generated whole transcriptome sequencing (WTS) on skin samples and blood-derived pDCs from ADEH^+^ (n=15), ADEH^-^ (n=13), and HC participants (n=13). First, exploring mechanisms related to dysregulated HSV-1 viral responses, we examined WTS from patient pDCs stimulated with mock- or HSV-1 infection. We then modeled pDC gene expression as a function of HSV-1 treatment, testing the interaction between infection and ADEH^+^ status. Although we found 9,079 differentially expressed genes (DEGs) in pDCs by HSV-1 infection (Table E1), there were no significant interactions with ADEH status. We also performed WGCNA co-expression analysis on the pDC samples, finding that 19 of 23 pDC co-expression networks were associated with HSV-1 infection (Fig. E1; Table E2). However, similar to the single-gene DE analysis, HSV-1-associated changes in expression of these 19 networks were not significantly different based on ADEH status (Fig. E1). In summary, we found no evidence that pDC response to HSV-1 infection is unique among ADEH^+^ patients.

We next explored whether ADEH^+^ skin exhibits gene expression changes, performing WTS on non-lesional epidermal and dermal skin (separated from skin biopsies) of the same ADEH^+^, ADEH^-^, and HC participants (Fig. E2). To examine this, we performed single-gene DE analysis between ADEH^+^ and HC participants, finding 14 ADEH^+^ DEGs in the dermis (Fig. 1A; Table E3). Upregulated dermal DEGs included known AD genes: *CCL18* chemokine, inflammatory tissue protease, *MMP12*, and fibroblast growth factor, *FGFR3*. Novel dermal DEGs included collagen (*COL6A6)* and cytokeratin (*KRT2*) genes, and the serine protease inhibitor, *SERPINB12*. Interestingly, we found two genomically adjacent (11p11) genes, troponin subunit I2 (*TNNI2*) and synaptotagmin 8 (*SYT8*), both upregulated in ADEH^+^ participants. Comparing epidermal expression in ADEH^+^ vs. HC, we observed 129 DEGs (Fig. 1A; Table E3), many of which have known roles in innate and adaptive immunity, inflammation, and cytotoxic immune functions, as well as contain multiple epidermal differentiation complex (EDC) genes and genes involved in epidermal growth and maintenance (Fig. 1A).

**FIG 1:**
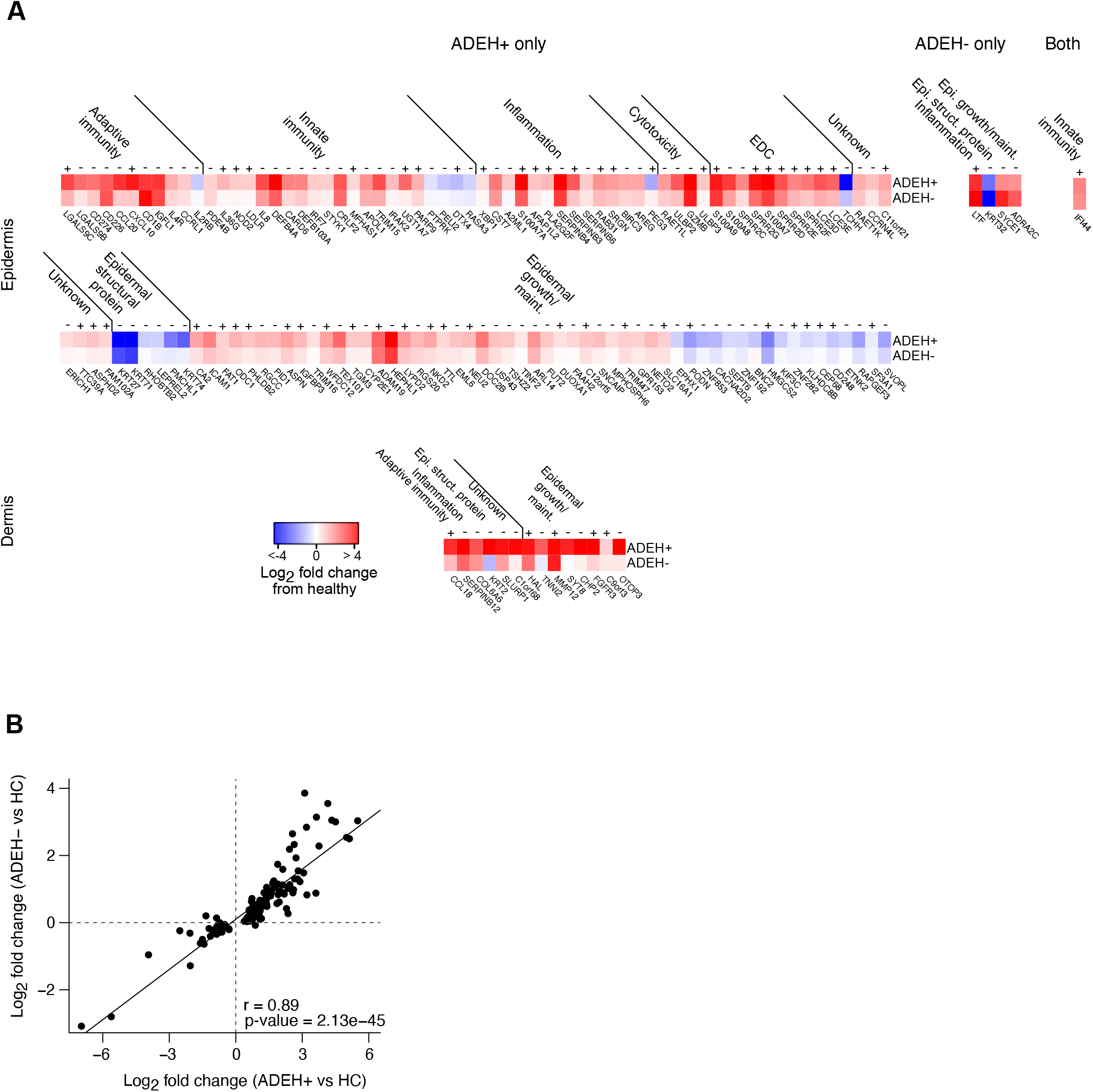
Multi-tissue transcriptome analysis of non-lesional skin from AD subjects with and without EH reveals dysregulated epidermal expression among subjects. **A**, Heat maps showing log_2_ fold change (LFC) in expression between ADEH^+^ (top row) or ADEH^-^ (bottom row) compared to HC for genes differentially expressed in one or the other disease group or both (from left to right), in epidermis (top) or dermis (bottom). The genes are organized into broad functional groups. **B**, Scatter plot comparing log fold changes between ADEH^-^ and HC epidermal skin with those between ADEH^+^ and HC epidermal skin, based on genes with significantly modified expression in ADEH^+^ epidermis compared to HC. Pearson correlation between the two sets of LFCs is indicated and demonstrates that genes dysregulated in ADEH^+^ skin are similarly dysregulated in ADEH^-^ individuals.

In contrast, when comparing skin gene expression of ADEH^-^ to HC participants, we observed no significant DEGs in dermal tissue and only five significant DEGs in epidermis, where only one epidermal DEG (*IFI44*) was shared between ADEH^+^ and ADEH^-^ groups (Fig. 1A; Table E3). However, when comparing fold changes in expression of ADEH^+^ DEGs between ADEH^+^-vs-HC and ADEH^-^-vs-HC analyses, the direction of effect (up- or down-regulation) for the ADEH^+^ and ADEH^-^ analyses was concordant for 98% and 86% of the epidermal and dermal DEGs, respectively. Furthermore, there was a strong positive correlation between ADEH^+^ DEG fold changes and fold changes observed for these genes in the ADEH^-^ group (Pearson r=0.89, p=2.13e-45, Fig. 1B), with consistently less pronounced dysregulation observed among ADEH^-^ participants. To further explore whether ADEH^+^ pathobiology is shared with ADEH^-^ patients, we examined the overlap between our ADEH^+^ DEGs and AD genes reported in five published RNA-seq studies of AD^14, 20-23^. Here, we found that the majority of ADEH^+^ epidermal DEGs (55/129 or 43% for non-lesional AD skin and 111/129 or 86% for lesional AD skin) and dermal DEGs (5/14 or 36% for non-lesional skin and 12/14 or 86% for lesional AD skin) have been previously associated with AD in at least one of the five studies. Together these results suggest that skin pathobiology underlying AD complicated by EH is similar in kind, but more severe in form, than that exhibited by AD patients without EH history.

### Dysregulation of EDC gene expression in the skin of ADEH^+^ patients corresponds with T2 inflammation

Examining the functional basis of ADEH^+^ differential expression observed in the epidermis, we found strongly increased expression of terminal skin differentiation pathways. Specifically, we found 10 EDC genes with significantly higher expression in the ADEH^+^ group, including five among the top ten ADEH^+^ DEGs (based on FDR). These upregulated EDC genes included five small-proline rich proteins (*SPRR2C, SPRR2D, SPRR2E, SPRR2F, SPRR2G*), two late-cornified envelope proteins (*LCE3D, LCE3E)*, and three S100A-type proteins (*S100A7, S100A8, S100A9*; Fig. 2A). We also found strong downregulation of the epidermal structural gene, trichohyalin (*TCHH*), among ADEH^+^ subjects. While none of these 10 EDC DEGs exhibited a discordant direction of effect between ADEH^+^ and ADEH^-^ DE comparisons, the magnitude of dysregulation was consistently lower, though not significantly, in the ADEH^-^ group (Fig. 2A).

**FIG 2:**
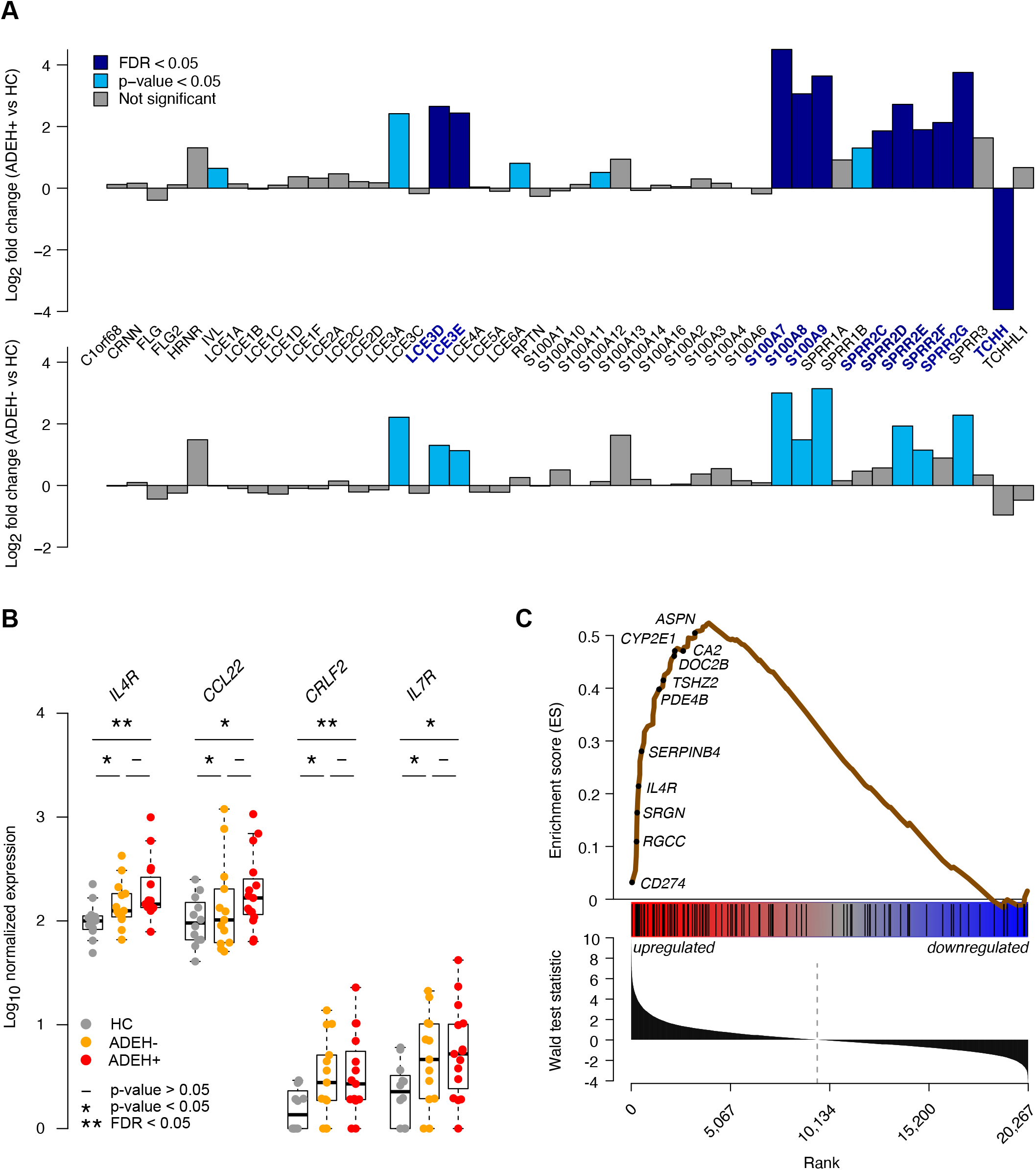
Dysregulation of EDC gene expression in ADEH^+^ corresponds with T2 inflammation. **A**, Bar plots of LFCs for EDC genes, comparing epidermal ADEH^+^ vs HC skin (top) and ADEH^-^ vs HC skin (bottom). Teal = nominally significant differences (p-value<0.05); dark blue = significant differences (FDR<0.05). Many dysregulated EDC genes in ADEH^+^ skin are similarly dysregulated in ADEH^-^ skin. **B**, Box plots of epidermal expression for canonical T2 genes in HC (grey), ADEH^-^ (orange), and ADEH^+^ (red) samples. “*” = nominally significant differences (p-value <0.05); “**” = significant differences (FDR<0.05). **C**, Enrichment score (ES) plot from GSEA visualizing enrichment of ADEH^+^ epidermal DEGs within a gene set pre-ranked based on DE analysis between T2-high and T2-low AD. *Top*: running-sum ES profile (brown line) across T2-ranked genes, where the left-side peak of positive ES values indicates particular enrichment of ADEH^+^ DEGs among the most T2-upregulated genes. A subset of genes belonging to the leading-edge of enrichment (appearing at/before the ES maximum and most strongly contributing to enrichment) are indicated. *Middle*: vertical black bars denote positions of ADEH^+^ DEGs along the T2-ranked list of genes. *Bottom*: bar plot showing the ordered distribution of the gene ranking statistic (Wald test statistic from DESeq2).

Potentially relevant to this upregulation of EDC genes is our previous work showing that T2 inflammation, particularly IL-4/IL-13 stimulation of keratinocytes, can elicit characteristic gene signatures in the skin, including dysregulated expression of EDC genes^24^. Supporting this, we found that two of the strongest skin biomarkers of T2 inflammation, *IL4R* and *CCL22*, were both upregulated in the epidermis of ADEH^+^ participants (Fig. 2B). Additionally, we found that both gene subunits of the TSLP receptor (*CRLF2* and *IL7R*) were significantly or nearly significantly upregulated in the dermis and epidermis of ADEH^+^ participants (Fig. 2B). To further explore ADEH^+^ upregulation in T2 inflammation, we used GSEA to test whether the 99 upregulated ADEH^+^ DEGs were enriched among genes previously found to characterize T2-high AD subjects^23^ (Table E4), finding strong enrichment of ADEH^+^ DEGs among genes most upregulated in T2-high subjects (p<0.001; Fig. 2C). Notably, eight of the 10 EDC genes upregulated with ADEH^+^ disease were among the leading-edge genes responsible for the T2-high enrichment. Together, these findings show that T2 inflammation is elevated among ADEH^+^ participants as a group and that this increased inflammation may contribute to observed dysregulation in EDC expression in the skin of ADEH^+^ patients.

### ADEH^+^ skin exhibits enhanced interferon and IL-36γ inflammation

Further examination of ADEH^+^-upregulated DEGs revealed multiple genes known to be upregulated by the epidermis in response to viral infection or interferon stimulation (Fig. 3A). Included among these was the interferon regulatory transcription factor 7 (*IRF7*), which drives expression of type I interferons and other viral response genes, and three members of the 2’-5’-oligoadenylate synthetase gene family (OAS), *OAS1, OAS2*, and *OASL*, which are interferon-inducible antiviral and innate immunity genes (Fig. 3A). To specifically address whether ADEH^+^ epidermal DEGs were more enriched in viral infection response genes, we leveraged published data identifying the transcriptomic response of keratinocytes to the viral mimetic, Poly(I:C)^25^ (Table E4). GSEA revealed strong enrichment of ADEH^+^-upregulated epidermal DEGs among the genes most induced by Poly(I:C) treatment of keratinocytes (p<0.001; Fig. 3B), including *CXCL10, ICAM1, IFI44*, and *IRF7*. Overall, these results support significant antiviral/interferon inflammation among ADEH^+^ patients.

**FIG 3:**
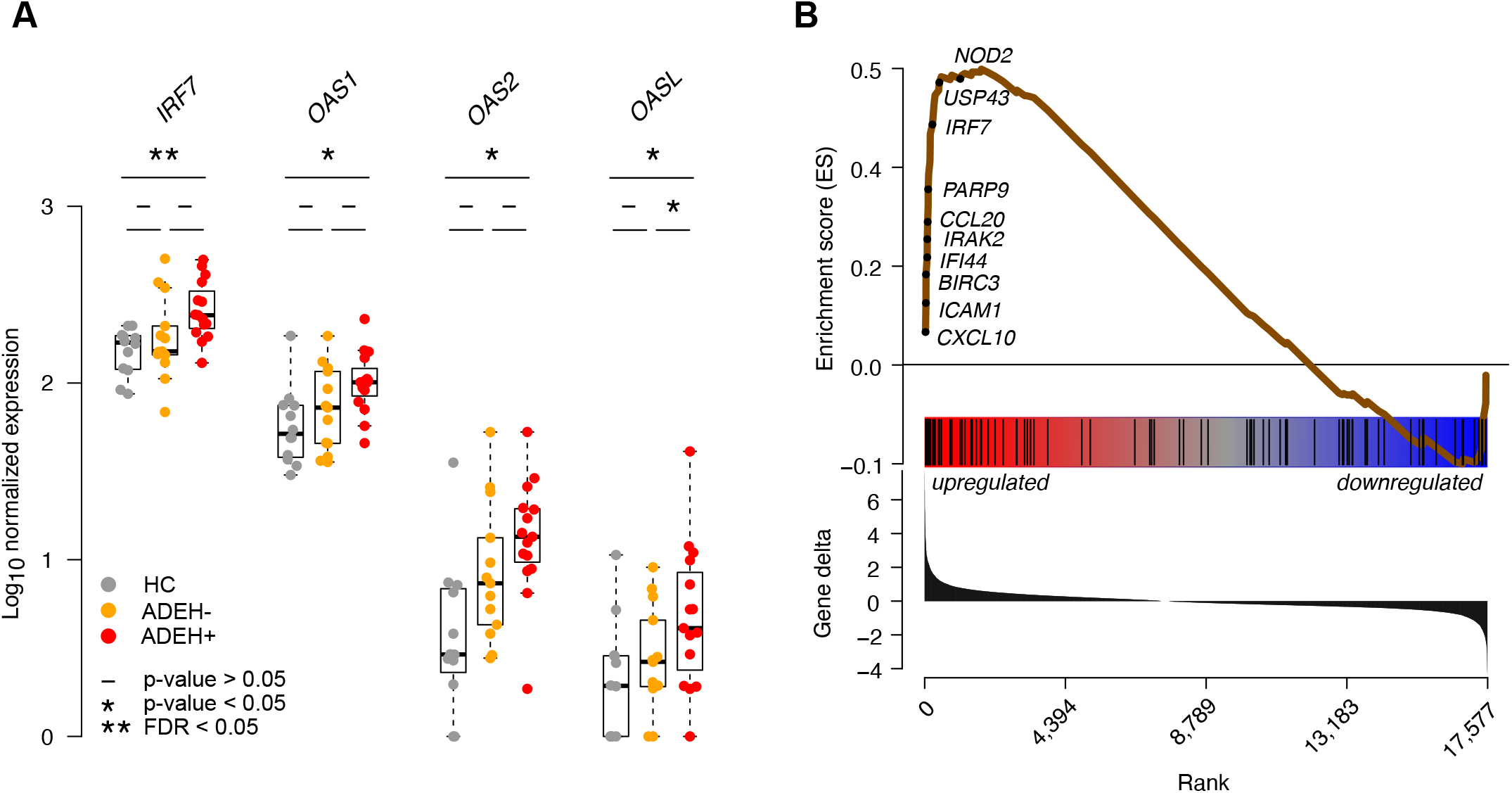
ADEH skin exhibits enhanced viral/interferon inflammation. **A**, Box plots of epidermal expression for canonical viral response genes in HC (grey), ADEH^-^ (orange), and ADEH^+^ (red). “*” = nominally significant differences (p-value<0.05); “**” = significant differences (FDR<0.05). **B**, Enrichment score (ES) plot from GSEA visualizing enrichment of ADEH^+^ epidermal DEGs within a gene set pre-ranked based on expression within poly(I:C)-stimulated keratinocytes compared to controls. *Top*: running-sum ES profile (brown line) across the poly(I:C)-ranked genes. A subset of leading-edge genes is shown. *Middle*: positions of ADEH^+^ DEGs along the poly(I:C)-ranked gene list. *Bottom*: ordered distribution of the gene ranking statistic (gene deltas).

Additionally, ADEH^+^ patients exhibited upregulation in expression of the inflammatory cytokine gene, interleukin 36γ (*IL36G*, FDR=0.01; Fig. 4A). IL-36γ is a member of the interleukin-1 family and has been reported to be prominently upregulated in psoriatic skin, but also in some AD cohorts, where it can drive expression of other chemokines and disrupt skin cornification. To capture the transcriptional consequences of IL-36γ stimulation on the epidermis, we used an *in vitro* NHEK culture system to differentiate keratinocytes in the presence and absence of IL-36γ protein. Whole transcriptome RNA-seq analyses revealed that IL-36γ treatment altered expression of 5,596 genes (FDR<0.05, Table E4) in differentiated keratinocytes. Of 129 genes modified in ADEH^+^ epidermal skin, over half (53%) were among these IL-36γ-stimulated DEGs and log fold changes associated with ADEH^+^ disease and IL-36γ stimulation were significantly correlated (p=2.02e-5, r=0.38, Fig. 4B). Remarkably, among the genes strongly upregulated in both datasets were all 10 of the ADEH^+^-upregulated EDC genes, (*LCE3D/E, SPRR2C/D/E/F/G*, and *S100A7/8/9)*. Moreover, based on GSEA, we found strong enrichment of the set of ADEH^+^-upregulated DEGs among the genes most induced by IL-36γ treatment of keratinocytes (p<0.001, Fig. 4C). All 10 ADEH^+^-upregulated EDC genes were among the 33 leading-edge genes identified in the IL-36γ enrichment analysis (Fig. 4C). The only EDC gene that was downregulated among ADEH^+^ subjects, *TCHH*, was also strongly downregulated by IL-36γ treatment (FDR=1.36e-34; Fig. 4B). Taken together, these results strongly suggest that both interferon and IL-36γ-driven inflammation are elevated in the epidermis of ADEH^+^ subjects.

**FIG 4:**
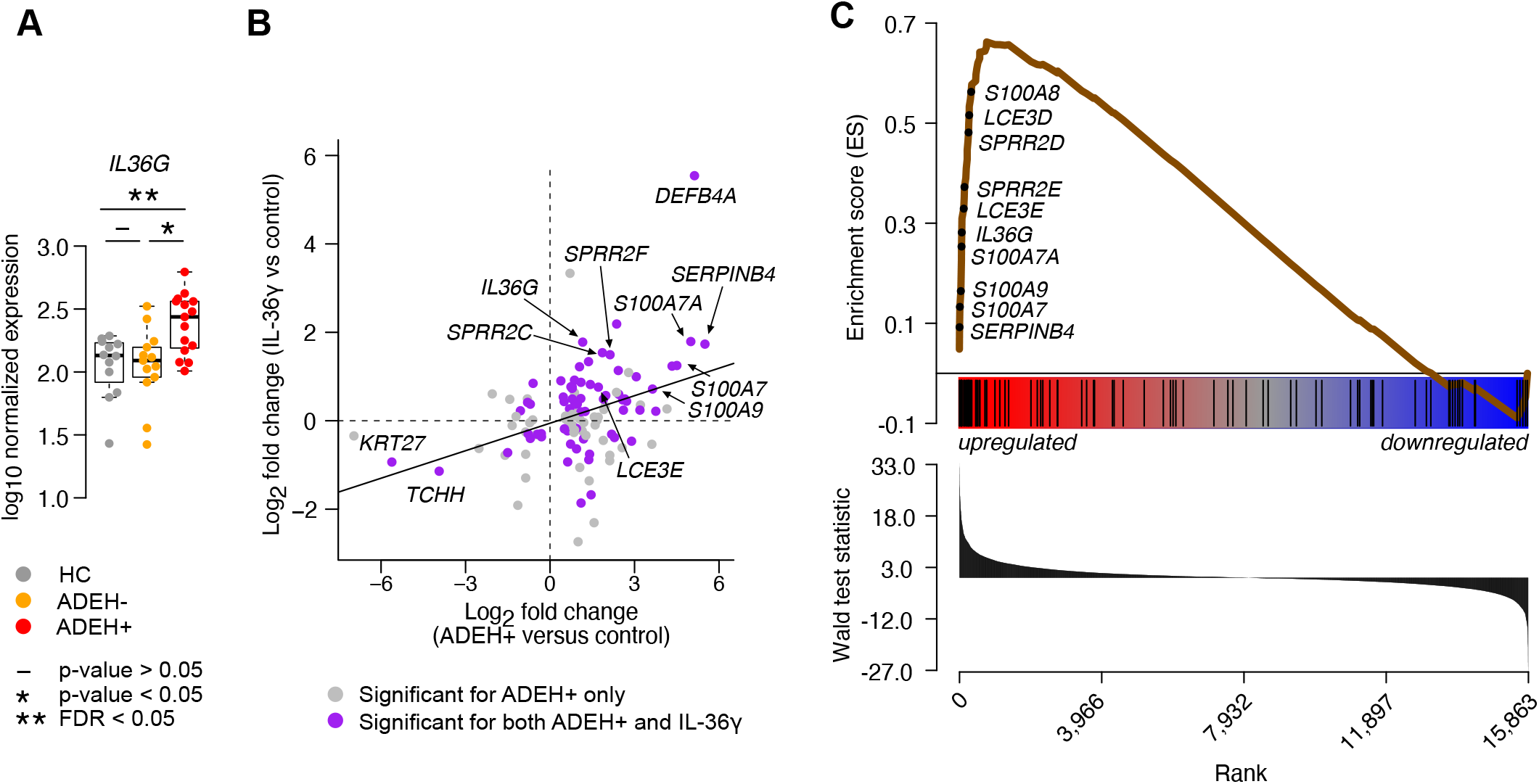
ADEH skin exhibits enhanced IL-36γ driven inflammation. **A**, Box plots of epidermal expression for *IL36G* in HC (grey), ADEH^-^ (orange), and ADEH^+^ (red). “*” = nominally significant differences (p-value<0.05); “**” = significant differences (FDR<0.05). **B**, Scatter plot comparing LFCs for epidermal ADEH^+^ vs HC DEGs (x-axis) with LFCs of these same genes based on comparing IL36γ-stimulated keratinocytes to controls (y-axis). Gray = DEG only in the ADEH^+^ vs HC comparison; purple = DEG in both comparisons. **C**, Enrichment score (ES) plot from GSEA visualizing enrichment of ADEH^+^ epidermal DEGs within a gene set pre-ranked based on IL-36γ-stimulated keratinocytes vs controls. *Top*: running-sum ES profile (brown line) across the IL-36γ-ranked genes. A subset of leading-edge genes is shown. *Middle*: positions of ADEH^+^ DEGs along the IL-36γ-ranked gene list. *Bottom*: ordered distribution of the gene ranking statistic (Wald test statistic from DESeq2).

### ADEH^+^ disease is characterized by co-expression of multiple inflammatory skin endotypes

We next investigated the shared genic aspects of the epidermal T2, interferon, and IL-36γ inflammatory expression signatures, finding that 39% of signature genes were within two or more of the three inflammatory programs (Fig. 5A). Notably, the T2/IL-36γ overlapping genes included eight of the 10 ADEH^+^-upregulated EDC genes. Despite this shared inflammatory character, we also identified 9, 13, and 23 genes specific to the IL-36γ, interferon, and T2 inflammatory programs, respectively (Fig. 5A).

**FIG 5:**
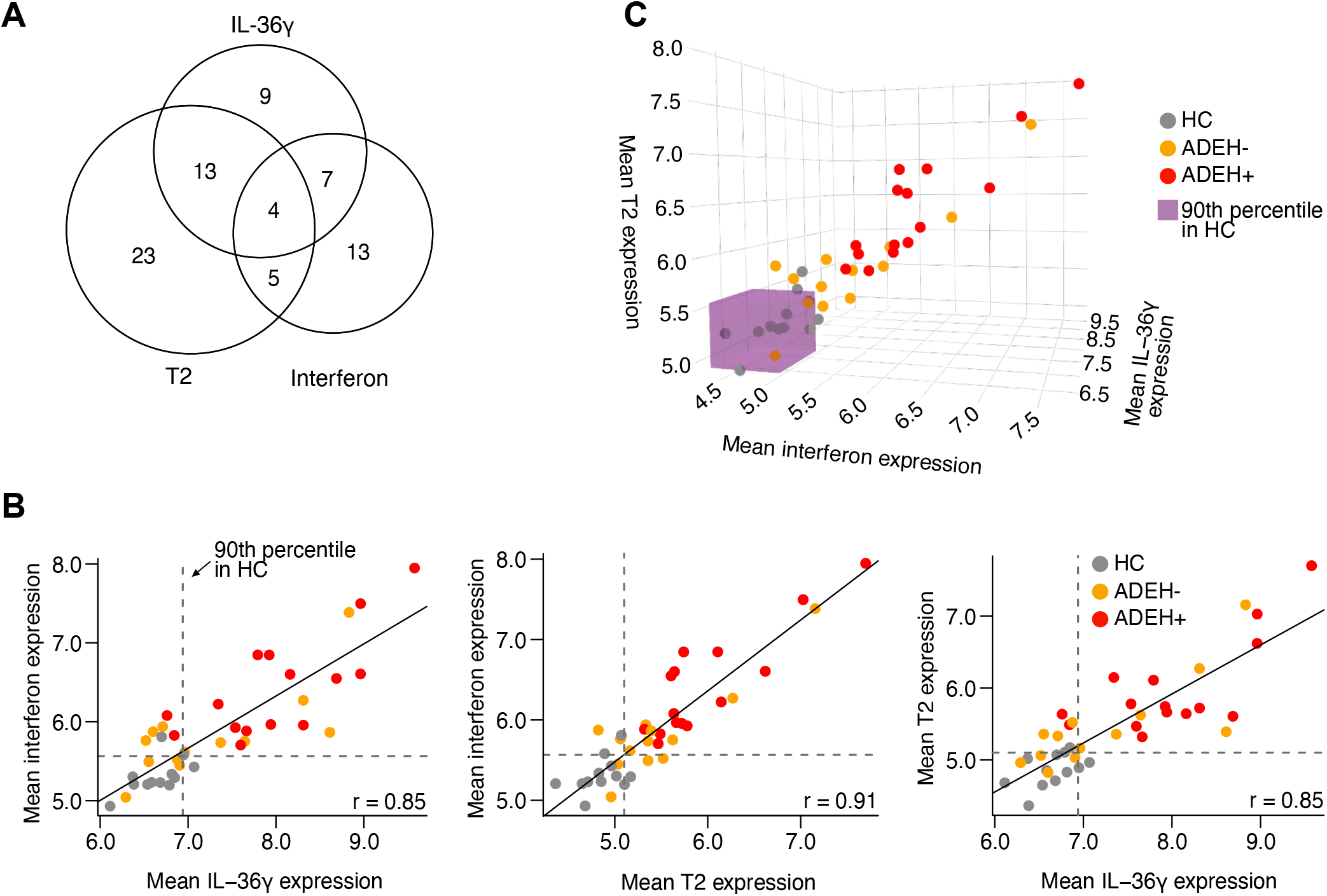
ADEH subjects co-express multiple inflammatory skin endotypes. **A**, Venn-diagram of leading-edge genes detected in distinct GSEAs that test for enrichment of epidermal ADEH^+^ DEGs within pre-ranked genes based on T2-high, viral-induced (poly(I:C)-stimulation), and IL-36γ-stimulated inflammation. **B**, Pairwise scatter plots of mean expression for distinct leading-edge genes from the three inflammatory endotypes (T2, viral, and IL-36γ). Grey = HC, orange = ADEH^-^, red = ADEH^+^; dashed lines = the 90^th^ percentiles of mean expression for HC samples; solid line = slope of the linear relationship; r = Pearson correlation coefficient. **C**, 3D scatter plot of mean expression for distinct leading-edge genes from the three inflammatory endotypes. Point colors are as in **B**; the purple box subsumes all values that are below the 90^th^ percentile threshold of mean expression for HCs.

Although T2, interferon, and IL-36γ inflammatory signatures were upregulated on a group level among ADEH^+^ patients, it is unclear whether this upregulation is uniform across all ADEH^+^ patients or driven by a subset of participants with high expression levels (i.e., an inflammatory endotype group). Supporting this possibility, prior studies have found significant inflammatory heterogeneity across the AD population.^1, 10-14^ Therefore, we leveraged mean expression of the T2, interferon, and IL-36γ-specific gene sets to dichotomously classify or “endotype” all participants for these inflammation patterns, defining participants as “inflammatory endotype-positive” or “negative”, based on whether their mean expression levels were greater or less than the 90^th^ percentile of the metric among healthy controls, respectively. Strikingly, we found that all 15 ADEH^+^ subjects were positive for the T2 and interferon endotypes, while 87% were positive for the IL-36γ endotype (Fig. 5B). In contrast, presence of these endotypes was much more variable among ADEH^-^ patients, with 69%, 69%, and 46% of this group positive for the T2, interferon, and IL-36γ endotypes, respectively (Fig. 5B). These patterns show that multiple inflammatory endotypes co-occur simultaneously in non-lesional AD skin, particularly among ADEH^+^ patients, where all three endotypes are nearly universally present, potentially underlying the more severe AD disease among ADEH^+^ patients. Moreover, we found that expression levels for the pairwise combinations of inflammatory metrics were all strongly positively correlated (Fig. 5B, C; r≥ 0.85), further suggesting a strong relationship among these different inflammation types.

## DISCUSSION

Here, we considered whether the basis for ADEH^+^ disease pathobiology lies in unique transcriptional changes to the skin (epidermis and dermis) and/or to pDCs (an immune cell population critical to antiviral responses). We uncovered significant transcriptional dysregulation in the epidermis of ADEH^+^ patients. The ADEH^+^ disease profile, which included upregulation of EDC genes, was similar in form to that observed for ADEH^-^ patients (here and previously published^12, 14, 20-22^), but was more intense in degree. These alterations in EDC expression were accompanied by epidermal inflammation consistent with stimulation by T2, interferon, and IL-36γ cytokines. Each of these inflammatory endotypes was observed among subgroups of ADEH-patients; however, ADEH^+^ patients were unique in their propensity to exhibit either two or all three of these inflammatory endotypes simultaneously. Together our findings suggest that ADEH^+^ disease is characterized by a complex, multi-faceted inflammation of the epidermis, accompanied by a dysregulated epidermal skin layer.

AD has traditionally been known as a T2-associated disease^26, 27^, which has been supported in many studies.^12, 28, 29^ However, recent genomic profiling of skin samples from AD patients has revealed significant heterogeneity in disease mechanisms, where T2-dominated inflammation is only observed in a subset of AD patients.^10-13, 30^ Here, we found only a subgroup of ADEH^-^ patients were T2-high, whereas all 15 ADEH^+^ patients were classified as T2-high, exhibiting increased expression of the skin T2 biomarker genes, *IL4R* and *CCL22*, and co-receptor genes for TSLP (an epithelial T2 alarmin previously associated with AD^31^), *CRLF2* and *IL7R*. These findings suggest that excessive T2 inflammation may play a significant role in ADEH^+^ disease, possibly through dysregulation of the normal innate immune mechanisms that protect skin from viral infection. Supporting this, food allergic AD patients with T2 skin inflammation have been found to exhibit enhanced trans-epidermal water loss.^32^ In fact, IL-4 and IL-13 stimulation of the skin downregulates loricrin and involucrin expression, critical skin barrier genes, and increases risk of *S. aureus* infection.^33, 34^ Additionally, T2 cytokines decrease skin production of multiple antimicrobial peptides, including human β-defensin 2 & 3 and cathelicidin, LL37^35, 36^.

Here we found that eight of the ADEH^+^-upregulated DEGs were EDC genes, all previously shown to be upregulated among T2-high AD patients.^12^ These EDC genes (*LCE3D, LCE3E, SPRR2C, SPRR2D, SPRR2E, SPRR2F, S100A8* and *S100A9*) encode proteins that contribute to structural stability of the cornified envelope and function as skin AMPs. Late cornified envelope (LCE) genes have long been recognized to play a role in formation of a healthy skin barrier, as the proteins they encode are cross-linked into the cornified epithelium.^37^ The 18 human LCE genes are arranged into 6 groups based on sequence and expression similarity. LCE genes in groups 1, 2, 5, and 6 are expressed in normal skin^38^ whereas group 3 LCE genes, which we found to be upregulated among ADEH^+^ patients (LCE3D/3E), are induced by injury and are strongly upregulated in lesional psoriatic skin.^38^ Furthermore, multiple genetic studies have confirmed that LCE3B/3C deletion is associated with psoriasis risk.^39, 40^ Importantly, LCE proteins have recently been shown to exert potent antimicrobial activity, including against *S. aureus* and *P. aeruginosa*; the species selectivity and strength of this antimicrobial activity varies by LCE group.^37^ Further supporting this differential antimicrobial activity of LCE group proteins, patients with the LCE3B/3C genetic deletion have been found to exhibit a unique microbiome profile.^37^ Also, among the ADEH^+^-upregulated EDC genes were the inflammatory alarmin genes S100A7/A8; these proteins form a heterodimer called calprotectin that exerts significant antimicrobial skin activity through the sequestration of calcium.^41, 42^ Although these genes have been found to be upregulated in AD, they are better known as psoriasis biomarkers and to be induced by IL-17 signaling.^42, 43^ The third set of ADEH^+^-upregulated EDC genes, Small Proline-Rich Protein 2 (SPRR2) genes, are known to cross-link the cornified envelope of the skin,^44^ but more recently were found to act as AMPs through disrupting bacterial membranes.^45^ Moreover, mice lacking the Sprr2a gene were found to be more susceptible to *S. aureus* and *P. aeruginosa* infections.^45^ We hypothesize that the selective upregulation of these EDC genes may alter both the barrier function and skin microbiome of ADEH^+^ patients, rendering their epidermis more susceptible to viral infection.

We also observed strong upregulation of *IL36G* gene expression in the epidermis of ADEH^+^ patients, along with significant enrichment of ADEH^+^ DEGs for genes upregulated by IL-36γ stimulation of keratinocytes, including all eight of the ADEH^+^-upregulated EDC genes. IL-36γ belongs to the IL-1 cytokine superfamily and is strongly associated with development of psoriasis,^46^ being elevated in skin lesions of psoriasis patients^47^ and underlying Th17-driven mouse models of psoriasis.^48, 49^ Moreover, loss-of-function mutations in IL36RN (an IL-36 receptor antagonist) are strong risk factors for development of pustular psoriasis.^50-52^ In comparison with psoriasis, only a few studies have implicated IL-36γ inflammation in AD, showing increased IL-36γ in AD skin lesions.^53, 54^ One possible reason for elevated IL-36γ in AD relates to *S. aureus* infection, a well-known feature of AD^55-57^ that is also elevated in ADEH^+^ disease.^58^ Recent studies using topical application of *S. aureus* on the skin surface of IL36R-knockout mice demonstrated that IL36R-mediated signaling was essential to skin inflammatory responses,^48, 49^ suggesting that IL-36 cytokines may be critical in *S. aureus*-induced skin inflammation in ADEH^+^ patients. Supporting the profound influence of IL-36γ signaling on skin biology, we found that IL-36γ stimulation of keratinocytes led to dysregulated expression of thousands of genes, including those involved in antimicrobial defense and epidermal barrier function. Together, our findings support an important role for IL-36γ-driven inflammation in ADEH^+^ pathobiology.

Based on our previous observation that PBMCs isolated from ADEH^+^ subjects produced fewer type I and type II interferons upon HSV-1 stimulation *ex vivo*,^6, 9^ we hypothesized that we might also observe diminished interferon programming in (1) pDCs, one of the major type I and type III interferon-producing peripheral immune cells^59^ and/or (2) the epidermis/dermis of ADEH^+^ individuals. However, we observed no reduced interferon signaling among ADEH^+^ patients in either cell/tissue type, with or without HSV-1 stimulation. Surprisingly, we instead observed significantly increased interferon activation in the epidermis of ADEH^+^ vs ADEH^-^ patients. As a first line of defense against virus infection^60^, an enhanced interferon program would be expected to diminish risk of viral infections. However, in the context of it being co-activated with T2 and IL-36γ programs, skin barrier dysfunction brought on by this hyper-inflammatory state may outweigh any antiviral benefits of elevated interferon expression. In addition to the unknown effects of this interferon inflammation on skin function, it is also unclear by what mechanism this inflammation is provoked. We note that interferon inflammation has been observed in the epithelium of another atopic disease: the airway epithelium of a subset of asthmatics.^61^ In fact, T2 inflammation may itself promote interferon inflammation, as suggested by reports that interferon inflammation and cellular stress are induced in epithelial cells stimulated with IL-13.^62^ Similarly, although acute AD is most associated with elevated T2 inflammation, chronic AD may invoke an elevated interferon inflammatory profile.^63^ Another intriguing possibility is that recurrent HSV-1 skin infections experienced by ADEH^+^ patients may epigenetically orient the epithelium toward an interferon program, often referred to as inflammatory memory.^64^ For example, in the gut, inflammatory epithelial remodeling originally prompted by a single infection persists, even after the infection resolves.^65^ We speculate that such inflammatory memory could be encoded within skin stem cells, which then give rise to keratinocytes more prone to expressing the interferon endotype. This phenomenon was demonstrated in mice, where inflammation-challenged epithelial skin stem cells showed epigenetic memory of exposures by exhibiting enhanced responses to subsequent injury. ^66^ Moreover, dysregulated inflammatory memory resulting from prior exposure in ADEH^+^ patients specifically was recently supported by the observation of persistent long-term abnormalities in the sphingosine-1-phosphate signaling system in the skin and plasma of these patients.^67^ Such epigenetic imprinting may underlie the heightened inflammatory milieu of ADEH^+^ disease and calls for further study.

Despite these exciting findings, we note several limitations to this study. First, as a cross-sectional analysis performed in the context of established disease, we are unable to determine the sequence by which these inflammatory and epidermal profiles occurred over the course of ADEH^+^ disease development. Both longitudinal skin profiling studies of ADEH^+^ patients and mouse models of ADEH^+^ disease development are needed to delineate the stepwise ADEH pathogenesis process. Secondly, although our results here are significant, the cohort studied is limited in numbers; therefore, additional studies are needed to replicate our findings. Finally, although our studies implicate potential changes in skin barrier function and microbial dysbiosis underlying ADEH^+^ disease, mechanistic studies are needed to directly investigate whether these hypotheses are correct.

Despite these limitations, our study advances the understanding of ADEH^+^ disease pathobiology, uncovering a multi-faceted epidermal inflammation underlying this severe AD phenotype group, which is associated with dysregulated expression of cornified envelope genes critical to epidermal innate defense. This work should not only reorient ADEH^+^ research towards these pathways/genes, but also suggests that current or emerging inhibitor drugs directed against these pathways may be effective treatment options for this patient group.

## Abbreviations

AD: atopic dermatitis
ADEH^+/-^: atopic dermatitis with/without recurrent complication of eczema herpeticum
DE: differential expression
DEG: differentially expressed gene
EASI: Eczema Area and Severity Index
EDC: epidermal differentiation complex
EH: eczema herpeticum
FDR: false discovery rate
FLG: filaggrin
GSEA: gene set enrichment analysis
HC: healthy control
HSV: Herpes Simplex Virus
LFC: log2 fold change
MDS: multi-dimensional scaling
NHEK: Normal human embryonic keratinocyte
pDC: plasmacytoid dendritic cell
pfu: plaque forming unit
PBMC: peripheral blood mononuclear cell
SCORAD: Scoring Atopic Dermatitis
T2: type 2 cytokine
TMAP: torrent mapping alignment program
VST: variance stabilized transformation
WGCNA: weighted gene co-expression network analysis
WTS: whole transcriptome sequencing

