## Supplementary information for "Atopic dermatitis complicated by recurrent eczema herpeticum is characterized by multiple, concurrent epidermal inflammatory endotypes"

**SUPPLEMENTARY METHODS**

**Human Subjects Recruitment**

Non-lesional skin samples were obtained from 15 adult AD patients with history of EH (ADEH^+^) (6 male and 9 female patients; age [mean ± SD], 33.1 ± 10.5 years), 13 adult AD patients with no history of EH (ADEH^-^; 4 male and 9 female patients; age [mean ± SD], 37.7 ± 11.4 years) and 13 nonatopic healthy control (HC) subjects (3 male and 10 female subjects; age [mean ± SD], 41.0 ± 8.4 years) with neither a personal nor family history of atopy and no skin disease. At the time of skin sample collection, Eczema Area and Severity Index (EASI), Scoring Atopic Dermatitis (SCORAD) and Rajka-Langeland total scores were collected. In addition, for all study participants serum IgE levels were evaluated. All ADEH+ patients had prior history of ≥3 episodes of EH. All patients did not have active infection at the time of evaluation. All study subjects were HSV seropositive (per past documentation or current HSV test performed for this study). Skin cultures were examined for the presence of *S. aureus*. ADEH^+^ and ADEH^-^ groups were matched by AD severity. Study subjects’ characteristics are summarized in Table 1. Study subjects with AD did not receive topical corticosteroids, topical calcineurin inhibitors, or topical/oral antibiotics for 1 week before enrollment; however, use of topical corticosteroids and antibiotics was allowed in the HC group. Patients were not treated with systemic immunosuppressive medications for more than 1 month before enrollment in this study. The study was approved by the Western Institutional Review Board, WIRB protocol #20161695 and was conducted at National Jewish Health, Denver, CO. All subjects provided written informed consent before participation in the study.

**Skin Biopsy Collection and RNA Extraction**

2-mm skin punch biopsy specimens were collected from the non-lesional skin area of the upper extremities. The biopsy specimens were dissected into the epidermis and dermis after 1 hour of digestion at 37°C in 5 U/mL dispase diluted in PBS (catalog no. 354235; Corning, Corning, NY). Isolated epidermal and dermal samples were immediately submerged into RLT buffer with β-mercaptoethanol (Qiagen, Valencia, CA) and frozen at -80°C for future RNA isolation. RNeasy Micro Kits (Qiagen) were used according to the manufacturer’s protocol to isolate RNA from the epidermis and dermis.

**Peripheral blood pDCs isolation and HSV stimulation**

Heparinized venous peripheral blood was collected from each study participant. PBMCs were isolated using density gradient centrifugation in Ficoll^TM^. Immediately after purification, 5x10^7^ PBMCs were subjected to pDC-purification using EasySep^TM^ Human Plasmacytoid DC Isolation kit (STEMCELL^TM^ Technologies) according to the manufacturer’s instructions. pDCs were plated at 5x10^4^ cells/100 µL in complete RPMI 1640 medium (RPMI 1640, 10% fetal calf serum, supplemented with L-glutamine, HEPES and antibiotics) in 96-well plates, and mock- or HSV-1-treated (multiplicity of infection 1) for 20 hours. After treatment, cells were harvested for RNA extraction.

**Keratinocyte culture and stimulation**

Normal human embryonic keratinocytes (NHEK) were purchased from ThermoFisher Scientific and maintained in EpiLife medium containing 0.06 mM CaCl_2_ and S7 supplement in 5% CO_2_ at 37°C. For NHEK differentiation, cells were cultured in EpiLife medium containing 1.3 mM CaCl_2_ for 3 days, then treated with 200 ng/ml of IL-36γ (R&D systems) for 2 days. After treatment, the cells were harvested for RNA extraction. RNA-seq analysis and qRT-PCR were performed.

**RNA Transcriptome Gene Expression and Quality Control**

RNA-seq libraries were constructed and barcoded using the Ion AmpliSeq^TM^ Transcriptome Human Gene Expression Kit. Barcoded RNA-seq libraries were pooled and sequenced on the Ion Torrent Proton sequencer using P1 chips.

Sequencing reads were mapped to AmpliSeq transcriptome target regions with the torrent mapping alignment program (TMAP) and quantified with the Ion Torrent ampliSeqRNA plugin, using the uniquely mapping option. Duplicated sequences were removed from the FASTA file and incorrect amplicon locations were corrected as previously reported^1^. Two dermal baseline samples that had < 6 million assigned reads and one epidermal baseline sample that exhibited a clear dermal expression signature based on multi-dimensional scaling (MDS) analysis were removed from the dataset.

**Analysis of bulk RNA-seq data**

To compare gene expression among disease groups *in vivo*, we performed gene-level differential expression (DE) analysis using the *DESeq2* R package^2^. We adjusted p-values to control for false discovery rate (FDR) using the Benjamini-Hochberg method. Differentially expressed genes were those with FDR < 0.05. Long intergenic non-coding (LINC) genes and genes of uncertain function (LOC genes), as well as lowly expressed genes (not reaching at least 10 counts in three percent of samples) were excluded from analysis. To compare gene expression between paired pDC samples which were mock- or HSV-1 stimulated, we performed variance stabilized transformation (VST)-normalization of counts using *DESeq2* and then used the *lmerSeq* R package to run a linear mixed model that predicts VST expression as a function of treatment, including a random intercept for each subject and an interaction term between treatment and disease status to test whether response to treatment differs by disease status. To compare keratinocyte samples stimulated with or without IL-36γ, we used *DESeq2.* Gene set enrichment analyses (GSEA) throughout were performed using the EnrichR API^3^ (libraries: KEGG 2021, Reactome 2016, and GO 2021). Plots of gene expression were based on log_10_ counts normalized by size factor using *DESeq2*.

We carried out weighted gene co-expression network analysis (WGCNA) on *ex vivo* stimulated pDC samples to detect functional networks of co-expressed genes^4^ that may respond differently to HSV-1 based on disease status. For analysis, we used VST-normalized counts, excluding genes expressing less than 10 reads in 10% of samples. We ran WGCNA using the *WGCNA* R package^5^, based on signed Pearson correlations, a soft-thresholding power of 37, a minimum network size of 30, maxCoreScatter = 0.70, minGap = 0.30, and cutHeight = 1.

For GSEA, we ranked genes based on the Wald statistic as calculated using *DESeq2* for the T2 and IL-36γ-based rankings. We obtained these values from Dyjack et al. 2018^6^ for T2 and generated them ourselves for IL-36γ based on the transcriptomic responses of the IL-36γ-stimulated NHEK cultures. For rankings based on poly(I:C), we downloaded expression values for NHEKs treated with either poly(I:C) or vehicle from GEO^7^ (accession GSE92646) and then ranked genes based on shift in expression between vehicle and poly(I:C) (gene deltas). For each GSEA, we ran 1,000 permutations using GSEA v4.1.0^8^.

**SUPPLEMENTARY TABLE LEGENDS**

**Table E1:** Differentially expressed genes and pathways in pDCs stimulated with HSV-1. Output tables from lmerSeq, giving gene-specific differences in HSV-1-stimulated compared to mock-stimulated pDCs in HC (tab 1), ADEH^-^ (tab 2), ADEH^+^ (tab 3), and averaged across all three groups (tab 4). Tabs 5-7 give estimates for the interaction between HSV-1 stimulation and disease status. The last two tabs give results from EnrichR, with enriched terms and pathways of up (tab 8) and down-regulated (tab 9) genes based on the overall effect of virus across disease groups (tab 4). For each of the differential expression summary tables (tabs 1-7), contrast estimates, standard errors, degrees of freedom (df), t-test statistics, 95% confidence intervals, p-values, and adjusted p-values (FDR based on the Benjamini-Hochberg method) are given. For each of the enrichments in the enrichment tables (tabs 8-9), the originating enrichment library, enriched term, gene overlap (number of overlapping genes is before the underscore and number of genes in the pathway is after the underscore), p-value, adjusted p-value (Benjamini-Hochberg method), combined score (ln[p-value] * z-score), itemized overlapping genes are given.

**Table E2:** WGCNA network genes and their enriched pathways, based on blood pDCs stimulated and mock-stimulated with HSV-1. The first tab gives genes for each network and subsequent tabs give EnrichR results for each network. Enrichment tables are as described above for Table E1.

**Table E3:** Result tables for differential expression analysis in *DESeq2*, comparing pairwise disease groups (HC, ADEH^-^, and ADEH^+^) in epidermis (tabs 1-3) and dermis (tabs 4-6). Each table gives genes, mean expression in the baseline group (listed second in the tab title), log_2_ fold change, Wald statistic, p-value, and adjusted p-value (Benjamini-Hochberg method).

**Table E4:** Ranking of genes for GSEA based on differential expression analysis between T2-high and T2-low AD subjects (tab 1; Dyjack et al. 2018^6^), shifts in expression between keratinocytes stimulated with poly(I:C) versus vehicle (tab 2; Zhu and Garza 2017^7^), and differential expression analysis between keratinocytes stimulated or mock-stimulated with IL-36γ (tab 3; this study).

**SUPPLEMENTARY FIGURE LEGENDS**

**FIG E1: Similar responses to HSV-1 infection are observed in pDCs among healthy, ADEH^-^, and ADEH^+^ subjects.** Box plots of the change (delta) in WGCNA network expression (summarized by eigengene values) between paired *ex vivo* pDC samples treated with HSV-1 compared with mock treatment. A subset of networks that are significantly modified with treatment are shown and samples are stratified by disease status (healthy control, ADEH^-^, and ADEH^+^). There are no significant differences in network responses among the three disease groups.

**FIG E2: Broad overview of transcriptome variation within non-lesional dermal and epidermal skin from AD subjects with and without EH.** Multi-dimensional scaling (MDS) visualization of transcriptomic variation across *in vivo* dermal and epidermal samples. Point colors denote disease status (tan = healthy control, green = ADEH^-^, blue = ADEH^+^) and point shapes denote different tissue types (square = dermis, circle = epidermis). Enriched pathways for the most highly weighted genes for each axis extreme (low or high) of MDS dimensions 1 and 2 are given.

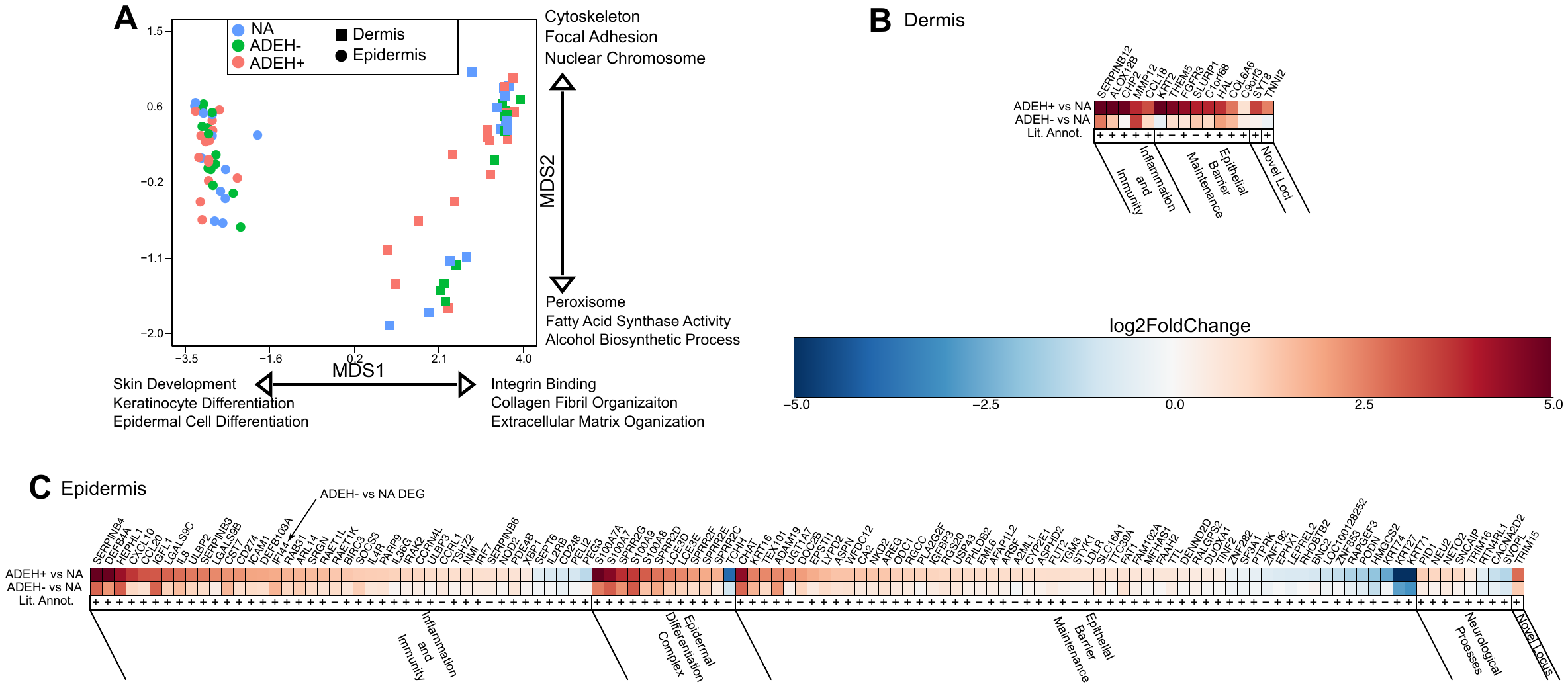

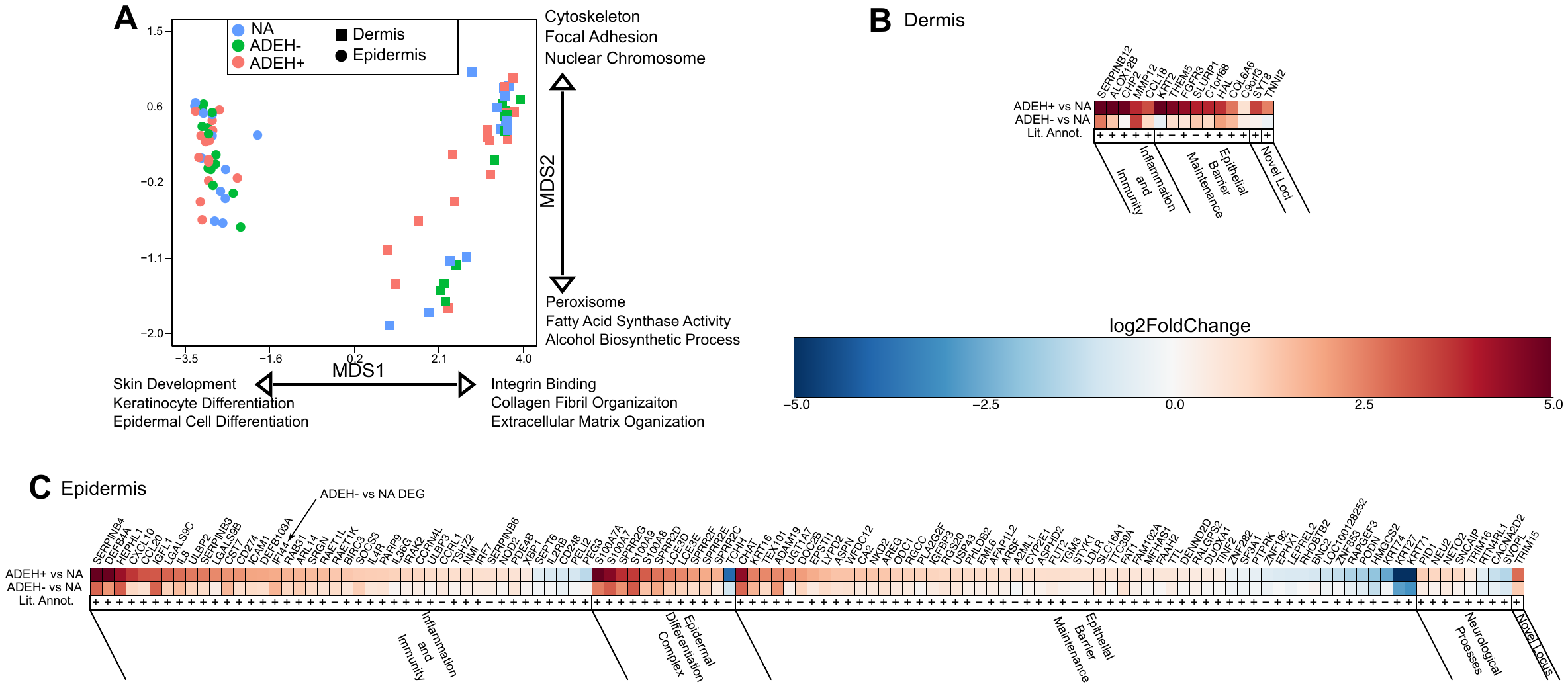
**REFERENCES**

1.	Poole A, Urbanek C, Eng C, Schageman J, Jacobson S, O'Connor BP, et al. Dissecting childhood asthma with nasal transcriptomics distinguishes subphenotypes of disease. Journal of Allergy and Clinical Immunology 2014; 133:670-8.

2.	Love MI, Huber W, Anders S. Moderated estimation of fold change and dispersion for RNA-seq data with DESeq2. Genome Biology 2014; 15.

3.	Chen EY, Tan CM, Lou Y, Duan Q, Wang Z, Meirelles GV, et al. Enrichr: interactive and collaborative HTML5 gene list enrichment analysis tool. BMC Bioinformatics 2013; 128.

4.	Zhang B, Horvath S. A general framework for weighted gene co-expression network analysis: The Berkeley Electronic Press; 2005.

5.	Langfelder P, Horvath S. WGCNA: an R package for weighted correlation network analysis. BMC Bioinformatics 2008; 9:559.

6.	Dyjack N, Goleva E, Rios C, Kim BE, Bin LH, Taylor P, et al. Minimally invasive skin tape strip RNA sequencing identifies novel characteristics of the type 2-high atopic dermatitis disease endotype. Journal of Allergy and Clinical Immunology 2018; 141:1298-309.

7.	Zhu AS, Li A, Ratliff TS, Melsom M, Garza LA. After skin wounding, noncoding dsRNA coordinates prostaglandins and Wnts to promote regeneration. Journal of Investigative Dermatology 2017; 137:1562-8.

8.	Mootha VK, Lindgren CM, Eriksson K-F, Subramanian A, Sihag S, Lehar J, et al. PGC-1α-responsive genes involved in oxidative phosphorylation are coordinately downregulated in human diabetes. Nature genetics 2003; 34:267-73.
