## Supplementary figures and images for "Atopic dermatitis complicated by recurrent eczema herpeticum is characterized by multiple, concurrent epidermal inflammatory endotypes"

### Supplementary Figure E1

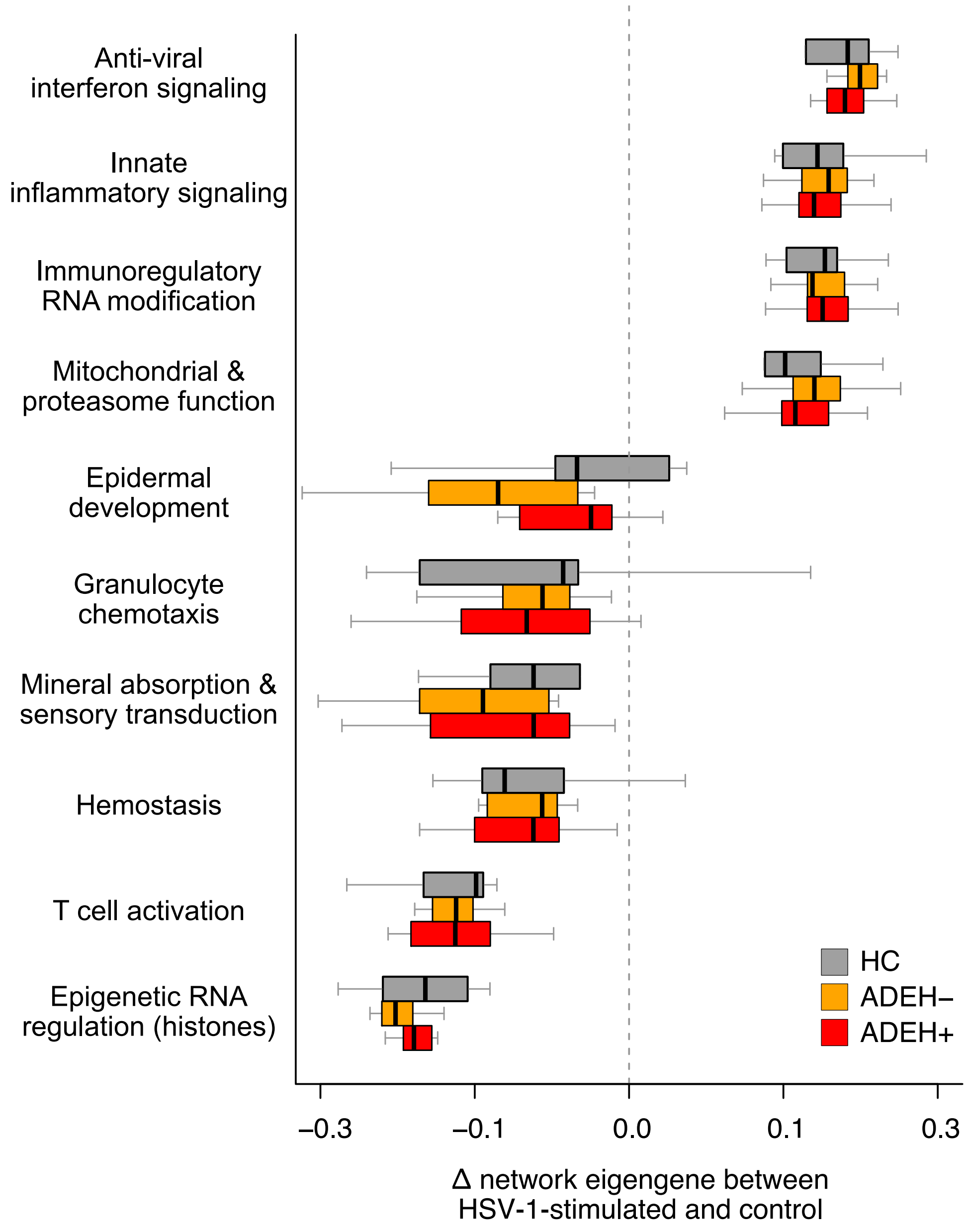

### Supplementary Figure E2

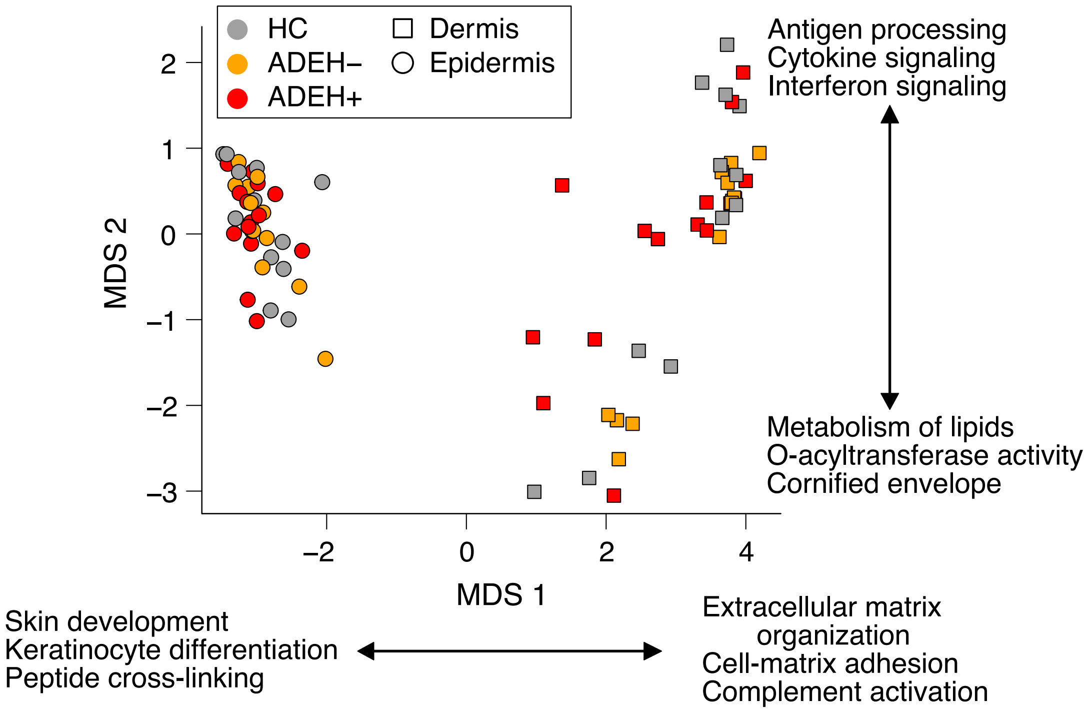
